# Platform Biological Divergence: quantifying gene-level differences between bulk and single-cell transcriptomics in breast cancer

**DOI:** 10.64898/2026.01.22.700982

**Authors:** Md Mamunur Rashid, Boon How Low, Kumar Selvarajoo

## Abstract

Bulk and single-cell RNA-seq offer complementary views of tissue transcriptomes. Nevertheless, pseudo-bulk (PB) profiles derived from single-cell data often differ systematically from true bulk measurements. These discrepancies are rarely quantified at the gene level, despite their impact on downstream analyses. To investigate the PB–bulk disagreement, here we introduce, based on linear mixture models, a gene-centric platform biological divergence (PBD) score. Across matched breast cancer, human adipose, and healthy retinal datasets, PBD stratifies genes into low-PBD “stable proxies” that retain true biological signals and high-PBD “divergent drivers” that shape PB–bulk differences. High-PBD (15–25%) genes show platform-specific signatures: bulk-enriched drivers highlight stromal, immune, and metabolic programs (breast cancer, adipose) and housekeeping/translational pathways (retina), whereas PB-enriched drivers reflect cell-intrinsic and regulatory processes, including immune, endothelial, progenitor, ECM, developmental, and photoreceptor programs. In contrast, low-PBD genes reliably preserve biology-driven variation and improve cross-platform concordance. Overall, the PBD framework reveals systematic gene-level biases between PB and bulk profiles, enabling more accurate comparison and integration of bulk and single-cell transcriptomes across tissues and disease contexts.

## 1. Introduction

High-throughput transcriptomics is transforming modern biology by providing multidimensional gene expression readouts at unprecedented resolution. Bulk RNA sequencing (RNA-seq) captures the average expression across cell populations, whereas single-cell RNA sequencing (scRNA-seq) profiles individual cells, revealing cell-to-cell variability[1–3]. This distinction is crucial for studying complex, heterogeneous diseases, such as cancer, where diverse cell types within the tumor microenvironment drive the pathophysiology. However, scRNA-seq comes with challenges, including high cost, technical noise, dropout events, and difficulty in detecting lowly expressed but biologically important genes, such as regulatory ncRNAs[4].

To mitigate these challenges, pseudo-bulk (PB) RNA-seq—aggregated from single-cell expression profiles—has emerged as a widely used surrogate for bulk RNA-seq in benchmarking deconvolution algorithms, differential expression (DE) analyses, cross-platform data integration, and synthetic data generation[5–9]. PB profiles reduce technical noise, improve statistical robustness, and facilitate downstream analyses[10]. Nevertheless, the fundamental assumption that PB faithfully reproduces bulk RNA-seq remains largely untested. Most prior evaluations focus on sample-level correlations, cluster concordance, or inferred cell-type composition, implicitly assuming that gene-specific differences can be corrected as batch effects[11–13]. This assumption is problematic because platform differences are not equivalent to batch effects. Batch effects reflect technical variability within the same measurement technology, whereas bulk and PB RNA-seq arise from fundamentally distinct data-generating processes[14,15]. In practice, platform-specific factors such as transcript capture efficiency, dropout patterns, cell-type imbalance, and amplification biases can introduce systematic gene-specific discrepancies that distort DE results, shift biological interpretations, and hinder cross-platform integration[16,17].

Current approaches, including correlation analysis, sparse principal component analysis (sPCA) to standard DE tools, and global distance metrics, are limited in their ability to detect gene-level divergence, particularly for genes whose expression differences are distribution-dependent rather than expression magnitude[18–21]. To address this gap, we introduce the **P**latform **B**iological **D**ivergence **(PBD)** score, a gene-centric, distribution-aware framework to quantify cross-platform divergence between PB and bulk RNA-seq. Applied to three independent human datasets—including breast cancer, adipose tissue, and retina samples—PBD stratifies genes into low-PBD stable proxies, whose PB expression preserves cross-platform biological and clinical signals, and high-PBD divergent drivers, dominate cross-platform inconsistencies undetectable by conventional DE or sparse PCA approaches. and enabling more reliable interpretation and integration of cross-platform transcriptomic data. Ablation analyses, further combined with gene ontology and cell-type–resolved expression mapping, identify biological processes most sensitive to platform effects, reveal the origins of PB–bulk divergence, and improve cross-platform interpretability.

## 2. Results

### 2.1 Overview of the cross-platform discrepancy analysis framework

The analytical workflow for identifying transcriptomic discrepancies between bulk RNA-seq and matched single-nucleus/single-cell–derived pseudo-bulk (PB) data is summarized in **Figure 1**. The pipeline consists of three key stages: 1) *Data acquisition and unified preprocessing*; Bulk RNA-seq and matched single-nucleus/single-cell datasets were acquired from the same samples, and PB RNA-seq profile we generated by aggregating cell-level counts. Both datasets were processed using a unified normalization, including low-variance gene filtering, library-size normalization, TMM adjustment, and log□ (CPM+1) transformation; 2) *Modeling platform-dependent discrepancies;* To identify platform-dependent gene sets, we applied conventional differential expression analysis and sparse PCA–based feature selection. We then compared these results with a variance decomposition (VD) framework based on a linear mixture model. Leveraging variance components estimated from the VD model, we defined a novel gene-centric **P**latform **B**iological **D**ivergence (**PBD**) score. The PBD scores displayed a clear bimodal distribution ranging from −1 to 1, enabling a data-driven cutoff (τ) defined at the local extrema of the density curve (Section 4.2); 3) *Gene categorization and downstream interpretation;* Genes with PBD > τ were defined as platform-divergence drivers (high-PBD), while genes with PBD < τ were defined as cross-platform stable proxy genes (low-PBD). Removing high-PBD genes enhanced bulk–PB similarity. Functional analysis of platform-divergent/high-PBD genes using gene ontology (GO) enrichment and cell–type–specific markers revealed key biological processes and cellular sources underlying cross-platform transcriptomic differences. Together, we hypothesized that major differences in gene expression profiles between matched PB and true bulk RNA-seq would be owing to biological factors, rather than purely technical artifacts.

**Figure 1.**
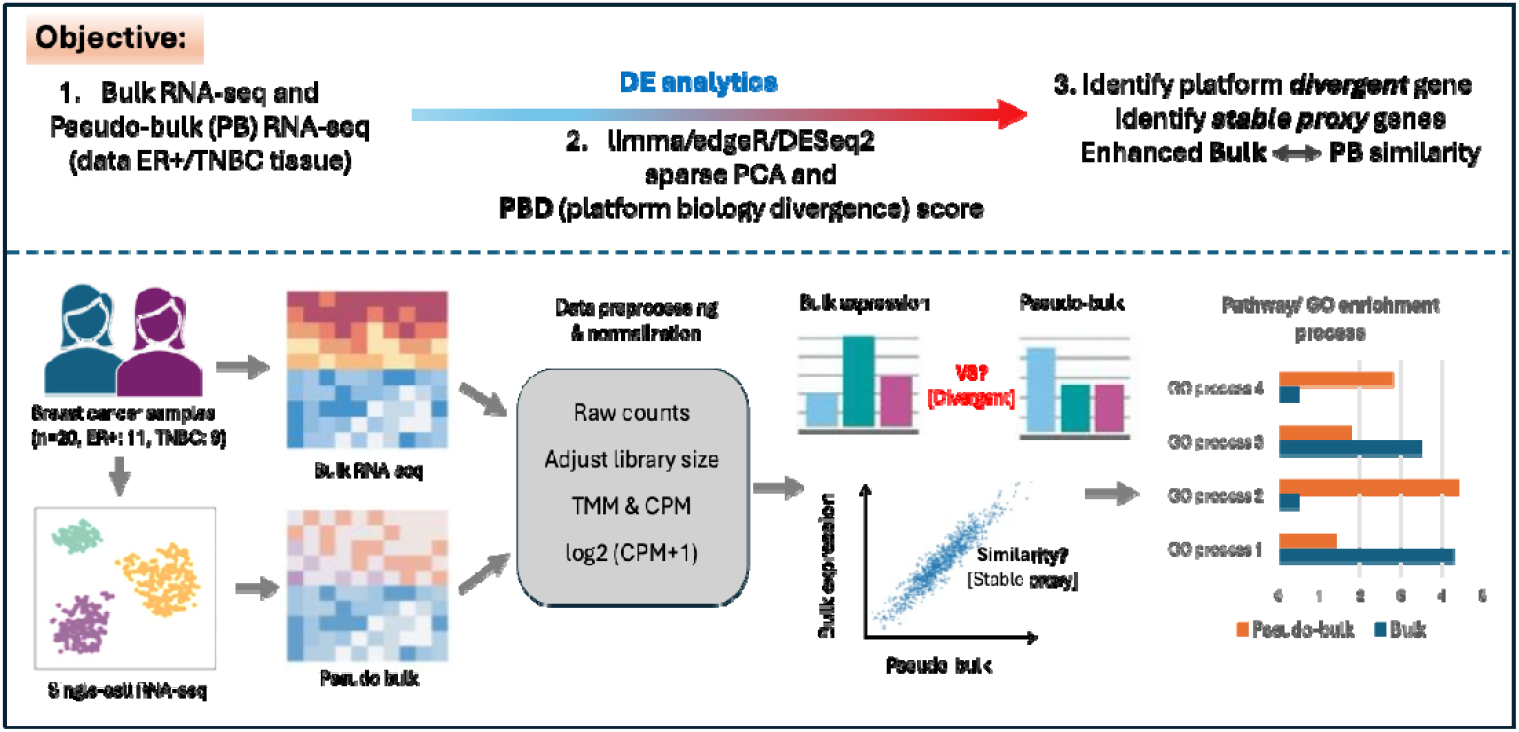
Workflow for assessing cross-platform biological discrepancies between bulk RNA-seq and matched pseudo-bulk RNA-seq. Paired bulk and single-cell/nucleus datasets were curated, and we simulated pseudo-bulk (PB) profiles by aggregating cell-level counts. Both datasets involved a unified preprocessing pipeline, including library size adjustment, between-sample TMM normalization, and log□ (CPM + 1) transformation. Gene-level discrepancies were detected using DE analytical approaches (above), including a variance decomposition–based linear mixture model, from which we derived a platform biology divergence (PBD) score for each gene. The bimodal distribution of PBD scores enabled data-driven classification of genes into cross-platform stable proxies and platform-divergent drivers. We then evaluated cross-platform similarity through correlation analyses, and assessed the biological relevance of platform-divergent drivers using GO enrichment and cell-type–specific marker analysis.

### 2.2 Biological discrepancies between bulk and pseudo-bulk RNA-seq

To characterize PB–bulk discrepancies, we first analyzed the single-cell landscape used to construct pseudo-bulk profiles (**Figure 2**). UMAP visualization of 40,858 single cells (subset of 68,102 available cells) identified nine major cell types, including cancer epithelial, normal epithelial, endothelial, cancer-associated fibroblasts (CAFs), perivascular-like (PVL) cells, and immune lineages (B cells, T cells, myeloid cells, and plasma blasts) (**Figure 2A**). This cellular composition forms the basis for PB aggregation and subsequent cross-platform comparisons.

**Figure 2.**
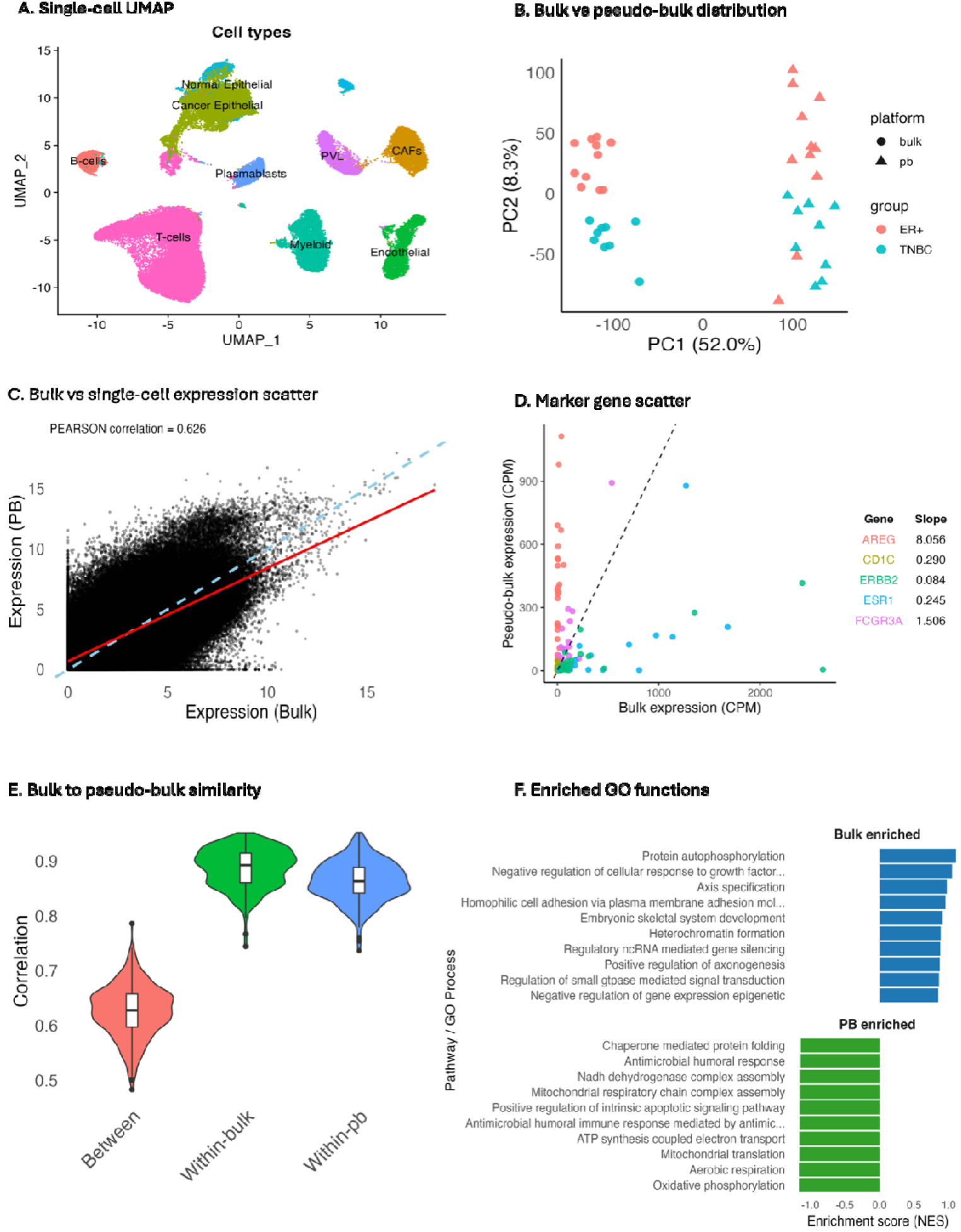
Systematic comparison of bulk and pseudo-bulk RNA-seq uncovers platform-dependent biological discrepancies across 20 breast cancer samples. (A) UMAP projection of single-cell transcriptomes showing the distribution of major cellular compositions used to construct pseudo-bulk profiles, including nine major cell types: T cells, B cells, myeloid cells, endothelial cells, perivascular-like (PVL) cells, cancer-associated fibroblasts (CAFs), normal epithelial cells, cancer epithelial cells, and plasmablasts. (B) Global expression distributions between bulk and PB RNA-seq (aggregated from single-cell data) from PCA analysis. (C) Gene-level scatter plot illustrating expression concordance and divergence between bulk and PB/single-cell RNA-seq. (D) Scatter distribution of representative marker genes, highlighting observed differences between bulk and single-cell profiles. (E) Correlation-based similarity analysis between bulk and PB gene expression profiles across samples. (F) Gene Ontology (GO) distinct biological processes enriched in bulk and PB transcriptomes, reflecting functional differences contributing to PB–bulk divergence.

**Figure 3.**
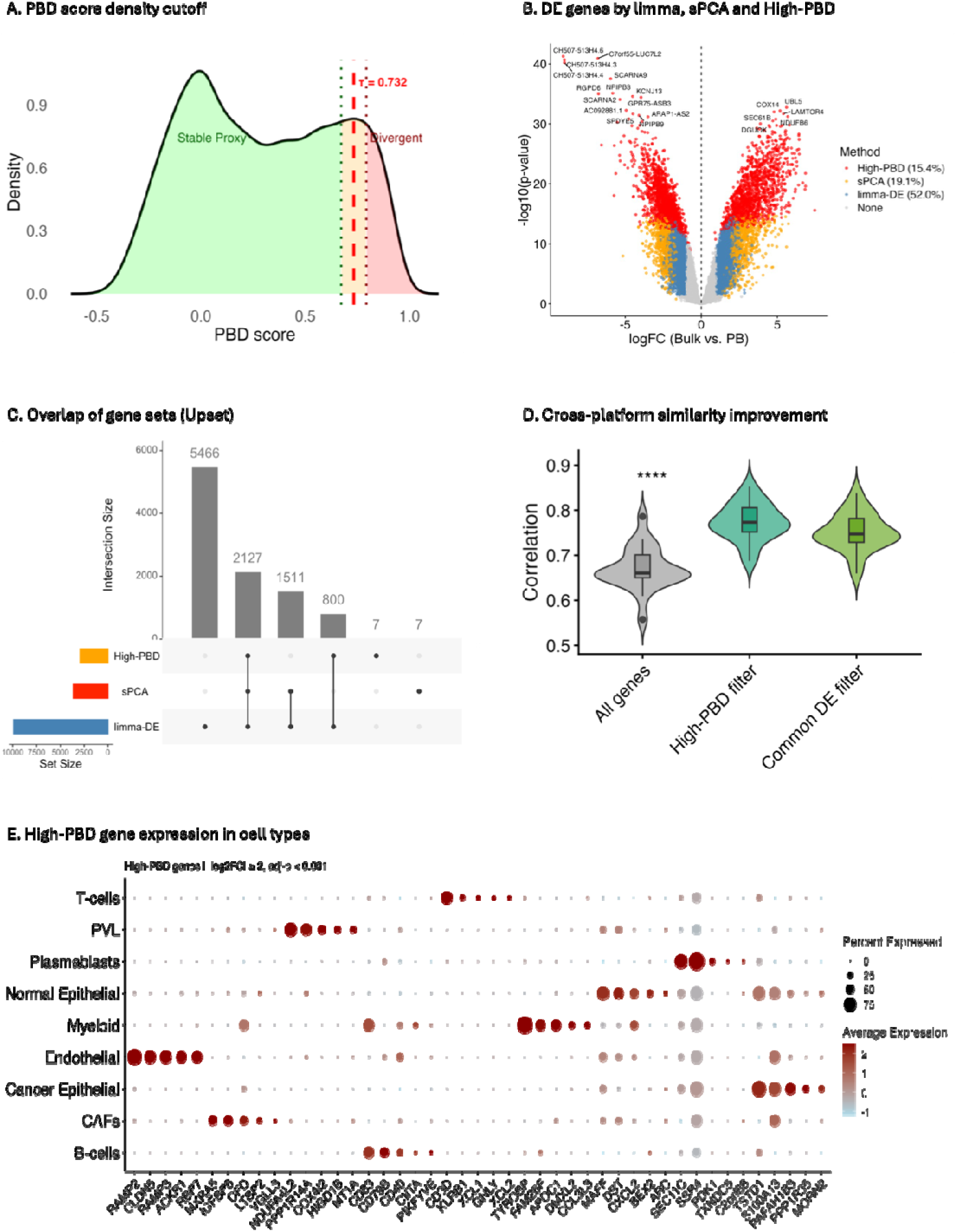
High-PBD genes reveal platform-dependent biology, and removing those enhances cross-platform similarity across 20 breast cancer samples. **(A)** Density distribution of platform biological divergence (PBD) scores, with the high-PBD gene cutoff window highlighted in red. Genes below this threshold represent cross-platform stable proxy genes, while genes above the threshold represent platform-divergent drivers. **(B)** Differentially expressed (DE) genes identified using *limma*, sPCA-selected features, and high-PBD gene prioritization. **(C)** UpSet plot showing the overlap and unique contributions of DE genes, sPCA-selected genes, and high-PBD gene sets. **(D)** Improvement in bulk–pseudo-bulk cross-platform similarity after filtering for high-PBD genes and intersecting common gene sets. **(E)** Expression patterns of high-PBD genes across annotated single-cell populations, highlighting cell–type–specific biological signals that drive cross-platform discrepancies

We next compared global expression patterns between bulk and PB samples. Principal component analysis (PCA) revealed a strong platform-driven separation along PC1; PB samples consistently exhibited positive PC1 scores, whereas bulk samples clustered in the negative PC1 region, despite being derived from the same patients. Biological variation, including disease subtypes (ER□, TNBC), was captured along PC2 but was markedly smaller than the bulk–PB separation. This indicates meaningful biological signals are retained yet embedded with a dominant platform effect (**Figure 2B**).

Gene-level overall concordance was evaluated using bulk-PB scatter plots. We observed that most genes aligned along the diagonal; however, a widespread mean shift (read line) was evident between platforms (**Figure 2C**). Scatter analysis of representative marker genes further provided biologically interpretable discrepancies; canonical tumor markers ESR1 and ERBB2 were elevated in bulk data, whereas AREG was more highly expressed in PB (**Figure 2D**). Immune markers (CD1C, FCGR3A) displayed variable representation contingent upon platform and cell-type abundance, aligning with prior studies of bulk and PB[22]. These patterns indicate that PB–bulk divergence may arise from both cellular composition differences and intrinsic platform biases. Kernel density estimates (KDE) further confirmed that PB profiles contained a higher proportion of low-expression genes, while bulk displayed a right-shift in highly expressed genes (**Supplementary Figure S1A**), consistent with scRNA-seq dropout effects[17]. In addition, the distribution of per gene PB–bulk mean deviations exhibited a log-normal–like shape centered near zero but spanning a wide range [−5 to +5], underscoring genome-wide mean-shift effects (**Supplementary Figure S1B**).

Correlation analyses further quantified cross-platform similarity. Median gene-expression correlations between matched bulk and PB samples were moderate (Pearson *r* ≈ 0.62), while within-platform correlations were substantially higher (r ≈ 0.75–0.88), underscoring that PB provides a biologically informative but divergent approximation of true bulk transcriptomes (**Figure 2E**).

Finally, to assess biological drivers of cross-platform divergence, we performed gene ontology (GO) enrichment using sparse PC1-selected gene sets. Bulk RNA-seq genes were predominantly involved in extracellular matrix organization, stromal activation, and macrophage-mediated processes, reflecting broad tumor microenvironmental contributions captured in bulk tissue (**Figure 2F**)[23]. In contrast, PB RNA-seq genes reflected mitochondrial electron transport, oxidative phosphorylation, ATP synthesis, and antimicrobial immune pathways, indicating a stronger influence of high-respiring epithelial tumor cells and activated immune subsets that are well represented in single-cell–derived profiles but diluted in bulk measurements[24].

Together, these results highlight that cross-platform discrepancies arise primarily from biological composition differences, with bulk emphasizing stromal and macrophage-rich pathways, while PB emphasizes metabolically active epithelial and immune-intrinsic programs.

### 2.3 Platform biological divergence (PBD) score identifies cross-platform divergent and true stable proxy genes

To quantify gene-centric PB–bulk divergence, we developed a two-step statistical framework. First, we applied variance decomposition (VD) using linear mixed-effect models to partition gene expression variability into biological and technical components. “Platform” (bulk = 1, PB = 0) was modeled as a fixed effect, while sample-, tissue-, and patient-level factors (e.g., sex, subtype) were modeled as random effects. The platform variance component is defined as the targeted component for assessing cross-platform effects. Next, we defined our **P**latform **B**iological **D**ivergence **(PBD)** score by measuring the deviation of each gene’s platform-associated variance from the average of all remaining variance components, as detailed in section 4.2 (Methods and Materials). By definition, negative PBD values correspond to purely conserved biological signals, whereas positive PBD values indicate moderate to strong cross-platform discrepancies. PBD scores ranged from −1 to +1 and formed a bimodal density distribution, enabling data distribution-aware selection of highly divergent genes using a local-extrema cutoff τ (**Figure 2A**). Genes below τ define low-PBD **“stable proxy”** genes, whereas genes above τ represent high-PBD **“platform-divergent drivers.”** Notably, disease subtype markers such as *ESR1, ERBB2*, and *AREG* resided in the low-PBD region (**Supplementary Fig. S2A**), confirming that the cutoff preserved true subtype-associated signals.

We then benchmarked high-PBD genes against features selected by limma differential expression (DE) and sparse PCA (sPCA). limma DE identified ∼52% of genes (n = 9,928; |logFC| > 2, FDR < 0.001), sPCA selected ∼19% (n = 3,647), and high-PBD identified ∼16% (n = 2,934) genes (**Figure 2B**). UpSet analysis showed a shared core of 2,127 genes across all methods (**Figure 2C**). However, the limma DE set was disproportionately large, reflecting sensitivity to global PB–bulk mean shifts and inflation of false positives under platform imbalance. In contrast, sPCA and PBD yielded more compact, interpretable feature sets. Variance partitioning (**Supplementary Fig. S2B–D**) further confirmed that PBD isolates most platform-driven divergence, whereas limma and sPCA capture blended effects of disease subtype and platform bias signals.

To validate PBD-defined gene categories, we assessed their distributional impact by repeating PCA after selectively removing each group. **Supplementary Fig. S2E–J** shows that high-PBD genes primarily drive bulk–PB separation, while low-PBD genes preserve ER□/TNBC subtype structure, consistent with transcriptome-wide mean-shift patterns observed earlier (**Supplementary Fig. S1B**). Accordingly, removing platform-divergent genes substantially improved cross-platform similarity, increasing median PB–bulk correlation to r ≈ 0.62 to 0.76 and enhancing KDE concordance (**Figure 2D; Supplementary Fig. S3B, D**). In contrast, high-PBD genes remained strongly discordant (r ≈ 0.24) with clear distributional mismatches (**Supplementary Fig. S2E; S3A, C**).

To determine the biological relevance of high-PBD genes, we mapped their expression across single-cell clusters and performed GO enrichment using GSVA. High-PBD (cross-platform divergence) genes exhibited elevated expression across epithelial, stromal, and immune compartments (**Figure 2E; Supplementary Fig. S4A**), reflecting cell-type–structured biology, not random technical noise. GO analysis further separated high-PBD genes into bulk-enriched developmental/stromal programs and PB-enriched metabolic/mitochondrial programs (**Supplementary Fig. S3E**). Bulk-enriched pathways involved ncRNA processing, axonogenesis, BMP response, and small GTPase signaling processes linked to tumor microenvironmental signals captured in bulk tissue[25]. In contrast, PB-enriched pathways highlighted oxidative phosphorylation, ATP synthesis, respiratory chain assembly, and antimicrobial responses, reflecting cell-intrinsic mitochondrial and immune programs robustly measured in scRNA-seq[26]. Violin plots further confirmed cell-type specificity of high-PBD genes (**Supplementary Fig. S4B**), underscoring their compositional coherence.

### 2.4 Applying the PBD score to the human adipose tissue for identifying cross-platform discrepancies

To validate and generalize the cross-platform differences observed in breast cancer, we applied and benchmarked the PBD score using 12 human adipose samples comprising paired bulk RNA-seq and snRNA-seq profiles[27]. The dataset included subcutaneous (SC/hSAT; 37,357 nuclei from 5 samples) and visceral (VIS/hVAT; 64,729 nuclei from 7 samples) adipose tissues. UMAP visualization of 35,262 high-quality single nuclei from hVAT identified eleven major cell types: adipocytes, ASPC1.APC, ASPC2.APC, mesothelial cells, endothelial cells, lymphatic endothelial cells (Lymph_Endo), pericytes, macrophages, monocytes, NK.T cells, and pre-myeloid cells (**Figure 4A**). The hSAT dataset contained eight cell types: adipocytes, ASPC1.APC, endothelial cells, pericytes, macrophages, monocytes, mast cells, and NK.T cells. These annotated cell populations formed the basis for pseudo-bulk (PB) aggregation and subsequent PB–bulk comparisons.

**Figure 4.**
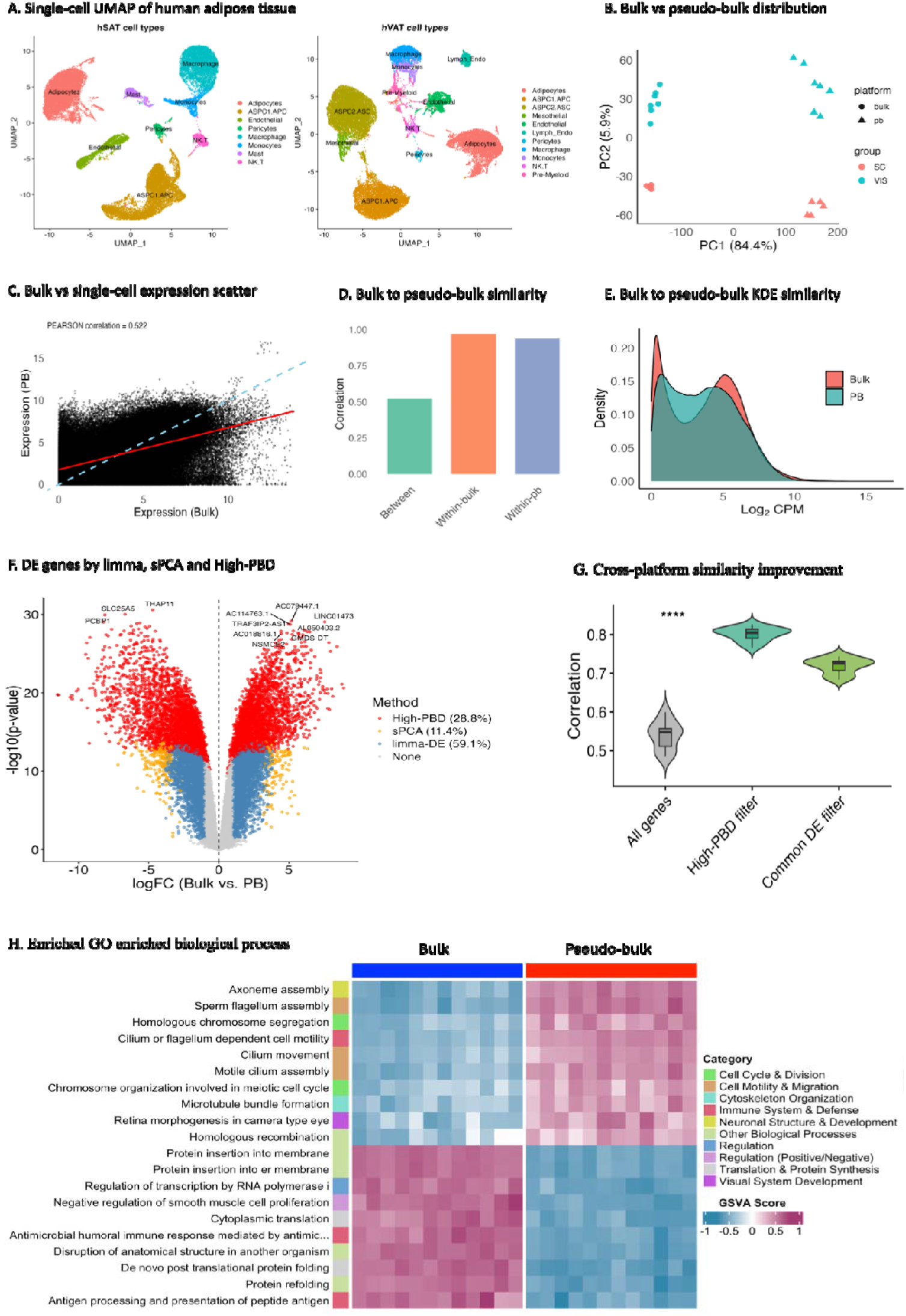
Quantifying cross-platform biology divergence (PBD) across 12 human adipose tissues using matched bulk and single-nucleus RNA-seq (snRNA-seq). **(A)** UMAP of snRNA-seq from subcutaneous (SC/hSAT; 37,357 nuclei) and visceral (VIS/hVAT; 64,729 nuclei) adipose tissue, showing 8 and 11 distinct clusters. **(B)** PCA of bulk and pseudo-bulk profiles separates samples by platform (PC1) and tissue type (PC2). **(C)** Gene-level scatter plot comparing bulk and pseudo-bulk expression, highlighting concordant and divergent genes. **(D)** Sample-wise correlation between bulk and pseudo-bulk gene expression matrices. **(E)** Kernel density estimation of global expression similarity, revealing overall concordance and systematic mean shifts. **(F)** Platform-divergent genes identified via differential expression (limma), sPCA, and high-PBD prioritization. **(G)** Enhanced bulk–pseudo-bulk alignment after removing high-PBD genes and intersecting divergent gene sets. **(H)** GO enrichment of biological processes enriched in bulk versus pseudo-bulk, highlighting functional pathways underlying transcriptomic divergence.

From PCA analysis, we observed a dominant platform-driven separation; PC1 explained 84.4% of the variance, with PB samples consistently occupying the positive PC1 region while bulk samples clustered negatively. Tissue subtype differences (SC vs. VIS) separated along PC2 (∼6% variance), indicating that while biological signals were retained, the platform effect remained the dominant source of variation (**Figure 4B**). Bulk and PB profiles displayed a larger mean expression shift (red line in **Figure 4C**), and their cross-platform concordance was moderate (Pearson r ≈ 0.52). In contrast, within-platform correlations were markedly higher (r ≈ 0.90–0.92; **Figure 4D**). KDE analysis further demonstrated that PB profiles contained a larger fraction of low-expression genes relative to bulk, which is consistent with expected snRNA-seq dropout effects (**Figure 4E**).

High-PBD genes identified in **Supplementary Fig. S5A–B** were benchmarked against gene sets selected by limma DE and sPCA. High-PBD identified ∼28.8% of genes (n = 5,586), limma DE selected ∼59.1% (n = 11,467; |logFC| > 2, FDR < 0.001), and sPCA selected ∼11.4% (n = 2,210) (**Figure 4F**). The disproportionately large limma DE gene set indicates inflation driven by global PB–bulk mean shifts rather than biologically meaningful differences. Variance partitioning across sample, tissue, and platform factors (**Supplementary Fig. S5C–E**) and PCA-based ablation analysis (**Supplementary Fig. S5F–K**) confirmed that high-PBD genes captured platform-specific variation, whereas low-PBD genes preserved SC/VIS tissue subtype structure. Removing high-PBD genes substantially improved cross-platform similarity, a median PB–bulk sample correlation increased r ≈0.79 (**Figure 4G**), KDE profiles became more concordant, and the bulk–PB expression distribution shift was significantly decreased (**Supplementary Fig. S6A**, right panels). In contrast, high-PBD genes remained highly discordant (r ≈ 0.08) and continued to exhibit prominent distributional mismatches (**Supplementary Fig. S6A**, left panels).

GO analysis of the adipose dataset showed that high-PBD genes in bulk-enriched terms were dominated by translation and protein-processing functions such as post-translational folding, cytoplasmic translation, ER/membrane insertion, and protein refolding, alongside antigen presentation and antimicrobial responses (**Figure 4H**). These reflect abundant housekeeping and immune transcripts that bulk sequencing captures efficiently. In contrast, PB-enriched pathways centered on cilium- and flagellum-dependent motility, axoneme and cilium assembly, microtubule bundle formation, and chromosomal segregation, representing structural and cell-cycle–linked programs better resolved in snRNA-seq–derived PB profiles. Mapping high-PBD gene further revealed cell–type–specific patterns (**Supplementary Fig. S6C**). Adipocytes showed the largest high-PBD set, including hallmark adipocyte regulators (ADIPOQ, FABP4), metabolic genes (ALDH2, ACADS), cytoskeletal and adhesion markers (ITGB1, MCAM), and numerous lncRNAs, indicating that bulk emphasizes abundant metabolic and secretory adipocyte programs, while PB captures nuclear, ciliary, and regulatory transcripts[28,29]. Conversely, ASPC1/APC progenitors were enriched for ECM and stromal remodeling genes (COL1A1/2, MMP2), fibroblast regulators (PDGFRB, ZEB1), and developmental signals (FGF14, ROBO2), reflecting the bulk’s greater sensitivity to stromal content[30,31]. Overall, bulk amplifies abundant metabolic and stromal programs, whereas PB highlights more specialized structural and cell regulatory features.

### 2.5 Applying the PBD score to quantify cross-platform discrepancy in human retinal benchmark samples

We next benchmarked the PBD score using a human retinal dataset of 20 healthy donors profiled with paired bulk RNA-seq and snRNA-seq[13]. Donors included 6 females and 14 males (ages 53–90), representing Asian (n=1), Caucasian (n=16), and Hispanic (n=3) backgrounds. After quality control, UMAP analysis of 34,856 high-quality nuclei (from 70,567 total) annotated six major retinal cell types: amacrine cells (AC), bipolar cells (BC), cones, horizontal cells (HC), non-neuronal/glial cells (NN), and rods (**Figure 5A**). These annotated populations were aggregated into pseudo-bulk (PB) profiles for PB–bulk comparisons.

**Figure 5.**
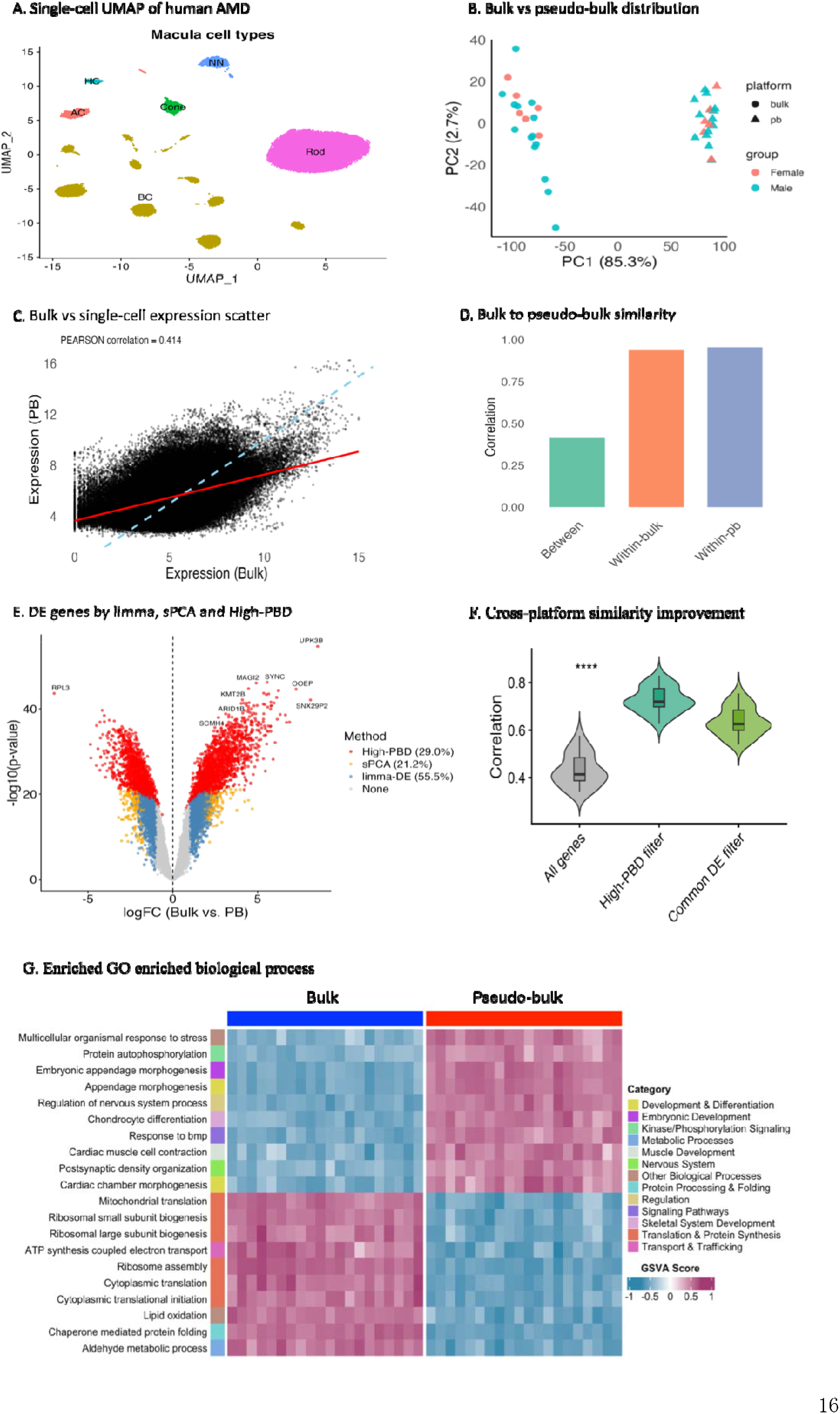
Quantifying platform biology discrepancies in 20 human retinal tissues using bulk RNA-seq and matched snRNA-seq. (A) UMAP embedding of snRNA-seq profiles from healthy (non-diseased) macular retina samples, showing six distinct cell clusters. (B) PCA of bulk and pseudo-bulk profiles separates samples by platform along PC1 and by individual variation along PC2. (C) Gene-level scatter plot comparing bulk and pseudo-bulk expression, highlighting overall similarity and platform-driven expression shifts. (D) Sample-wise correlation between bulk and pseudo-bulk gene expression matrices. (E) Platform-divergent genes identified using differential expression (limma), sPCA, and high-PBD prioritization. (F) Improved bulk–pseudo-bulk similarity after removing high-PBD genes and intersecting DE gene sets. (G) GO enrichment of biological processes enriched in bulk versus pseudo-bulk, revealing platform-specific biology underlying transcriptomic divergence.

PCA revealed a strong platform-driven separation along PC1 (85.3% of variance), with PB samples scoring positively and bulk samples negatively, while donor-level variation was retained on PC2 (∼2.7%) (**Figure 5B**). Cross-platform comparisons showed a larger mean expression shift (**Figure 5C**) and low concordance between PB and bulk (Pearson r ≈ 0.41), in contrast to high within-platform correlations (r ≈ 0.90; **Figure 5D**). KDE analysis further indicated that PB contained more low-expression genes, consistent with dropout effects (**Supplementary Fig. S7E**).

High-PBD genes were benchmarked against limma DE and sPCA-selected features. High-PBD identified ∼29% of genes (n=2,904), limma DE selected ∼55.5% (n=5,554; |logFC|>2, FDR<0.001), and sPCA selected ∼21.2% (n=2,118) (**Figure 5E**). The inflated limma gene set reflects global PB–bulk mean shifts sensitivity. Variance partitioning across donor, age, gender, ethnicity, and platform (**Supplementary Fig. S7A–D**) confirmed that high-PBD genes captured platform-specific variation, whereas DE and sPCA gene sets mixed platform effects with demographic components. The low-PBD genes markedly improved PB–bulk similarity, a median PB–bulk sample correlation increased to r ≈ 0.73 (**Figure 5F**), KDE distributions aligned more closely, and bulk–PB expression shifts were reduced (**Supplementary Fig. S7G, H**, right panel). In contrast, the high-PBD gene subset remained highly discordant (r ≈ 0.09) with persistent distributional mismatch (**Supplementary Fig. S7F, H**, left).

GO analysis showed that bulk-enriched high-PBD genes were dominated by core metabolic and housekeeping functions; ribosome assembly, protein folding, cytoplasmic and mitochondrial translation, aldehyde metabolism, and lipid oxidation (**Figure 5G**). This reflects abundant transcripts and biosynthetic processes captured in bulk RNA-seq[32]. PB-enriched pathways highlighted developmental and regulatory programs, including appendage and embryonic morphogenesis, nervous-system process regulation, BMP signaling response, chondrocyte differentiation, cardiac development, postsynaptic density organization, and protein autophosphorylation; low-abundance, signaling-linked features more readily resolved in snRNA-seq–derived PB profiles[16]. High-PBD genes also displayed clear cell-type specificity (**Supplementary Fig. S7I**). Rods contributed the largest high-PBD set (ABLIM3, KCTD8, NIPAL1), cones were enriched for ARR3, GNAT2, GRIK4, bipolar neurons for CA10, GABRA5, ZNF804B, glia for FOXP2, GABRG3, CNTNAP3B, and horizontal and amacrine cells for FRMPD4 and GRM8[33]. Overall, bulk emphasized abundant metabolic and housekeeping programs, whereas PB profiles highlighted finer regulatory, developmental, and cell-type–specific signaling features in the human retina.

## 3. Discussion

High-throughput transcriptomic analyses increasingly integrate bulk RNA-seq and single-cell RNA-seq (scRNA-seq) to capture complementary views of tissue biology. However, persistent discrepancies between these platforms complicate joint interpretation, limit model transferability, and pose risks when pseudo-bulk (PB) RNA-seq is used as a surrogate for bulk measurements in computational benchmarks, deconvolution, and other downstream analyses[6,7]. To address this, we developed the Platform Biological Divergence (PBD) score, a distribution-aware measure to quantify gene-centric cross-platform divergence and identify genes disproportionately influenced by platform-specific biology. Benchmarking PBD across three independent human datasets—breast cancer, adipose tissue, and retina samples—revealed unifying principles underlying the origins, magnitude, and biological basis of PB–bulk discrepancies.

First, across all datasets, PB and bulk profiles exhibited a dominant platform-driven axis of variation, often explaining >80% of total variance. This primary separation persisted even when donor, demographic, or biological covariates were controlled, indicating that technical and modality-dependent measurement properties outweigh inter-individual variability in shaping transcriptomic structure[15,17]. Consistently, PB profiles showed a higher proportion of very low-expression and dropout-prone genes, whereas bulk profiles were enriched for high-abundance metabolic and housekeeping transcripts[2,34]. These intrinsic differences between technologies directly contributed to global expression shifts and low PB–bulk correlation relative to within-platform similarity.

Second, the PBD score proved effective in isolating the subset of genes driving cross-platform divergence. Across the three datasets, high-PBD genes consistently captured platform-specific variation, while alternative selection strategies (e.g., limma DE or sparse PCA) were confounded by donor-level or biological effects. Importantly, removing high-PBD genes markedly improved PB–bulk similarity, reducing distributional mismatches, and harmonizing kernel-density profiles—demonstrating that a relatively small portion of platform-sensitive genes disproportionately drives global divergence.

Third, biological enrichment analyses showed that high-PBD genes were not purely technical artifacts but reflected platform-dependent biology. Bulk-enriched programs consistently highlighted extracellular matrix remodeling, stromal activation, macrophage signaling, and tissue-level metabolic pathways/processes dominated by abundant transcripts and cell types well represented in bulk tissue[25,35]. In contrast, PB-enriched signatures emphasized cell-intrinsic, low-abundance, or rare-cell programs such as T-cell activation, endothelial signaling, neuronal-specific pathways (retina), and mitochondrial metabolism linked to high-respiring epithelial or immune subsets[36]. These patterns reinforce the idea that PB and bulk emphasize different aspects of tissue biology rather than one modality being uniformly superior[14].

In summary, we present the Platform Biological Divergence (PBD) score, a robust gene-centric metric for quantifying systematic discrepancies between bulk RNA-seq and single-cell-derived pseudo-bulk transcriptome. Across three independent human datasets, PBD consistently identified platform-sensitive genes, revealed shared biological drivers of PB–bulk divergence, and demonstrated that removing high-PBD genes substantially improves cross-platform concordance. Bulk RNA-seq preferentially captured tissue-level stromal and microenvironmental programs, whereas pseudo-bulk profiles emphasized cell-intrinsic and rare-cell transcriptional signals. Collectively, the PBD score provides a principled framework for cross-platform gene filtering, integration, and interpretation, with broad applications in reference atlas construction, deconvolution benchmarking, differential expression analysis, and multimodal modeling.

### Limitations and future directions

While PBD effectively disentangles platform-dependent effects in matched datasets, its current formulation relies on paired bulk and single-cell sequencing measurements. Future extensions will focus on enabling unpaired cross-platform applications, allowing broader assessment of platform concordance, and facilitating integration across heterogeneous transcriptomic studies.

## 4. Methods and materials

### 4.1.1 Breast cancer cohort with clinical subtypes (GSE176078)

We obtained raw count data from the *Wu et al*. breast cancer cohort under the Gene Expression Omnibus (GEO) accession number GSE176078[37]. This dataset contains matched bulk and single-cell RNA-sequencing (scRNA-seq) profiles, along with clinical metadata from 20 breast cancer patients. Samples were annotated into three clinical subtypes: HER2-positive (HER2+, *n* = 4), estrogen receptor-positive (ER+, *n* = 11), and triple-negative breast cancer (TNBC, *n* = 9). To focus specifically on ER+ and TNBC biology, HER2+ samples were excluded, resulting in a final cohort of 20 patients for downstream analyses.

#### scRNA-seq processing and quality control

Raw scRNA-seq count matrices containing 100,064 cells and 29,733 genes were imported into Seurat to construct a Seurat object[38]. Cells were retained if expressed at least 200 genes, and genes were kept if detected in ≥10 cells. Mitochondrial transcript percentages were calculated per cell, and cells exhibiting extreme library sizes, feature counts, or mitochondrial proportions (defined as median ± 5 MADs) were removed. After filtering, 25,209 genes and 68,102 cells remained. To reduce computational load during visualization, ∼60% of cells from each major cell type were randomly subsampled while preserving proportional representation across samples. Filtered counts were log-normalized, and 3,000 highly variable genes were selected. Data were scaled, followed by principal component analysis (PCA). The first 20 PCs were used for UMAP visualization and clustering. To mitigate inter-sample batch effects, Harmony integration was applied before final UMAP projection and clustering. Cell type annotations were obtained directly from the pre-annotated metadata accompanying the dataset.

#### Bulk and pseudo-bulk RNA-seq processing

Pseudo-bulk (PB) RNA-seq profiles were generated by summing raw scRNA-seq counts across annotated cell types for each of the matched samples. To ensure a fair comparison, both PB and bulk datasets were processed using a unified pipeline. Genes with zero total counts across samples were removed, and a set of 19,057 common genes across all 20 donors was retained. Library size normalization for both platforms was performed using the TMM (trimmed mean of M-values) method implemented in edgeR (version 4.0.1) followed by log-transformation using log2(CPM + 1)[19,39]. Genes exhibiting zero variance in either the PB or bulk dataset were excluded before downstream statistical analyses.

### 4.1.2 Human adipose tissue benchmark datasets

We obtained paired bulk RNA-sequencing (RNA-seq) and single-nucleus RNA-sequencing (snRNA-seq) datasets from human visceral (hVAT) and subcutaneous (hSAT) adipose tissues[27,40]. Tissue samples were collected from individuals undergoing elective abdominal surgery, following written informed consent. The initial discovery cohort consisted of 7 hVAT and 5 hSAT samples.

For the single-cell component, snRNA-seq was performed on the same discovery cohort samples using the 10x Genomics platform, yielding 64,729 nuclei from hVAT and 37,357 nuclei from hSAT. Raw count matrices from both bulk and snRNA-seq were preprocessed and normalized following the procedures applied to the breast cancer datasets described above. UMAP visualization of the snRNA-seq data revealed 11 transcriptionally distinct clusters in hVAT and 8 clusters in hSAT, each corresponding to unique annotated cell types.

### 4.1.3 Human retinal benchmark dataset

We analyzed 20 healthy human retinal tissue samples collected within 6 hours postmortem using matched bulk RNA-seq and single-nucleus RNA-seq (snRNA-seq) profiling[13,41]. The snRNA-seq dataset contained 70,567 nuclei, spanning six major annotated cell types: amacrine cells (AC, 2,103 nuclei), bipolar cells (BC, 16,999 nuclei), cones (3,063 nuclei), horizontal cells (HC, 839 nuclei), non-neuronal/glial cells (NN, 3,254 nuclei), and rods (44,309 nuclei). Raw count matrices from both bulk and snRNA-seq were preprocessed and normalized following the procedures applied to the breast cancer datasets described above. Pseudobulk profiles were generated by summing UMI counts across all nuclei within each sample. The matched design ensures that bulk, snRNA-seq, and pseudobulk measurements share identical underlying cell-type compositions, enabling a controlled assessment of cross-platform discrepancies between conventional bulk RNA-seq and single-nucleus profiling.

### 4.2 Platform biological divergent (PBD) score gene analysis

To quantify systematic gene-level differences in expression distributions between bulk RNA-seq and scRNA-seq-derived pseudo-bulk (PB) profiles, we employed a two-step statistical framework based on variance decomposition and gene-wise divergence scoring. This approach explicitly accounts for the fact that bulk and single-cell technologies arise from distinct data-generating processes: bulk RNA-seq captures population-averaged expression, whereas single-cell RNA-seq reflects cell-to-cell heterogeneity aggregated during pseudo-bulk construction.

**First**, variance decomposition methods using linear mixed-effects models were used to split gene expression variance into biological and technical parts. For each gene g, the normalized expression across all samples is modeled as,

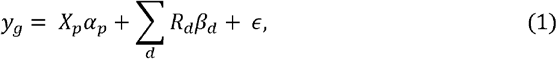

where *X*_*p*_ represents the design matrices for fixed effects *p*, including data sequencing platform (bulk=0, PB=1) with effect size 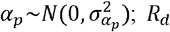 represents design matrices for random effects *d*, including donor, tissue, disease subtype, sex, and other biological covariates; *ϵ* and is the residual error term. Random effects were modeled as, 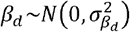and residuals as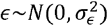.

Variance components were estimated via maximum likelihood as implemented in the “*variancePartition*” R package[42]. For each gene, the variance explained by random effects was calculated as,

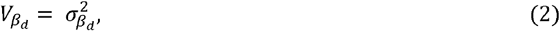

The platform fixed effect variance was defined as,

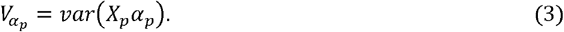

Residual variance was given by,

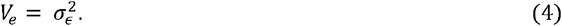

And the total variance for each gene was defined as,

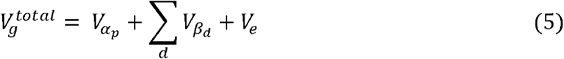

**Second**, to quantify relative platform-sensitive genes over preserved biological variations, we defined a gene-centric **platform biological divergence** (**PBD**) score as,

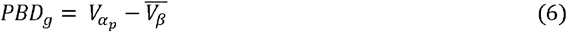

where 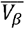 represents the mean variance proportion explained by random effect components, including donor, tissue, disease subtype, sex, and age. PBD scores are linearly scaled to the range [1,1], where negative values indicate genes dominated by conserved biological variation and positive values indicate genes primarily driven by platform-specific divergence.

A minimal threshold *τ* for identifying platform-sensitive genes was selected by analyzing the observed bimodal distribution of PBD scores, which reflects two distinct gene categories. The optimal separation point was defined as the last local maximum of the kernel density estimate, corresponding to the peak of the highly platform-sensitive gene subset. This data-driven strategy requires no parametric assumptions and yields biologically interpretable gene classification. Genes were classified as **divergent drivers** (PBD > τ) or **stable proxies** (PBD ≤ τ).

### 4.3 Principal component analysis (PCA) and sparse PCA

To assess global transcriptomic structure and visualize differences between pseudo-bulk (PB) and bulk RNA-seq samples, we performed dimensionality reduction using standard PCA and sparse PCA (sPCA). Log-transformed CPM values were 0-centered. sPCA was applied using the sparsepca::spca() function with α = 1e-4, and β = 1e-3 to obtain sparse loadings[18]. We focused on the first two components (PC1 and PC2), which captured the largest proportion of variance and were used to visualize platform separation. Non-zero sPCA loadings identified genes contributing most to platform-specific variation.

### 4.4 Cross-platform similarity analysis

To quantify transcriptomic similarity, cross-platform correlation was assessed using the Pearson correlation between all sample pairs, as calculated by the cor() function in R. The results were visualized using heatmaps and violin plots, which showed both within- and between-platform correlations.

### 4.5 Kernel Density Estimation (KDE)

To assess the observed data distributional shift, KDE was applied to log-transformed CPM data using *ggplot2*::geom_density(). This enabled a qualitative assessment of bulk-PB variations in gene expression distribution

### 4.6 Differential expression (DE) analysis

Differential expression analysis between PB and bulk RNA-seq profiles was performed using limma on log_2_CPM-transformed data[20]. Genes with |log□FC| > 2 and FDR < 0.001 were considered significant. Effect sizes from limma, sPCA loadings, and PBD values were jointly evaluated to identify genes with consistent platform-specific expression patterns. Volcano plots were used to visualize log□fold changes and FDR values, highlighting informative DE genes.

### 4.7 Gene set enrichment analysis

To characterize biological processes differentiating between bulk and PB transcriptomes, we applied Gene Set Variation Analysis (GSVA) to select DE gene subsets, using the GSVA R package (v1.x)[43]. The *MSigDB C5* Gene Ontology Biological Process (GOBP) collection was used as the reference database[44]. Sample-wise GSVA scores were computed for both platforms, and enriched differential pathway was assessed using two-sided *t*-tests with Benjamini–Hochberg correction (FDR < 0.05). Significant pathways were visualized using “*barplots*” and heatmaps from “*ComplexHeatmap*”[45].

### 4.8 Single-cell–level validation of platform-divergent genes

To validate high-PBD genes at single-cell resolution across major cell types, we performed differential expression analysis using Seurat’s *FindAllMarkers* function with the Wilcoxon rank-sum test. Genes were considered significant if they satisfied |log□FC| ≥ 2, adjusted *p* < 0.001, and a minimum detection fraction of ≥ 10%.

### 4.9 Visualization and statistical analysis

All analyses were conducted in R version 4.5.0. Visualizations included scatter plots, PCA and UMAP projections, correlation heatmaps, violin plots, volcano plots, Euler and UpSet diagrams, variance component plots, and single-cell expression plots (dot, feature, and violin plots)[46]. Detailed descriptions of statistical tests, parameters, and procedures are provided in the respective Methods subsections above.

## Declarations

### Ethics approval and consent to participate

Not applicable.

### Consent for publication

Not applicable.

### Data availability

Matched bulk RNA-seq and single-cell RNA-seq data, with clinical annotations were obtained from the Wu *et al*. breast cancer cohort under GEO accession GSE176078 [37]; Paired bulk RNA-seq and snRNA-seq datasets from human visceral (hVAT) and subcutaneous (hSAT) adipose tissues were obtained from previously published studies[27,40]; and matched bulk RNA-seq and snRNA-seq data from 20 healthy human retinal donors were obtained from published studies[13,41].

### Code availability

Source code for all analysis scripts and pipelines is available at GitHub (https://github.com/mamun41/PBD-Platform-Biological-Divergence-/tree/main)

### Author contributions

M.M.R. developed the computational pipelines for transcriptomic analyses, generated all visualizations, and wrote the manuscript. B.H.L. edited the manuscript and K.S. conceptualized the study, supervised, and edited the manuscript.

