## Supplemental Figures for "Platform Biological Divergence: quantifying gene-level differences between bulk and single-cell transcriptomics in breast cancer"

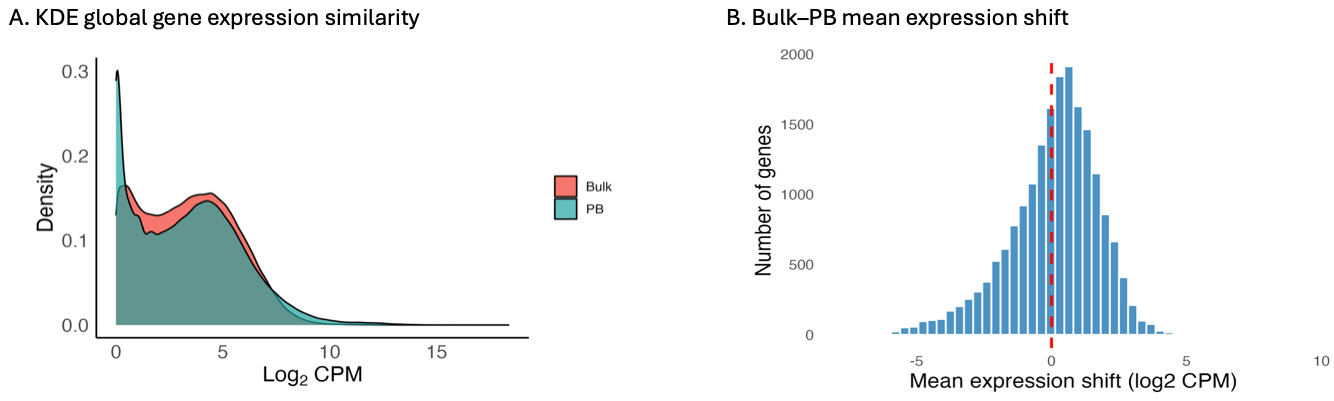

**Supplementary Figure** **S1. Global bulk–PB transcriptome comparison**

(**A**) Kernel density estimation (KDE) of global gene expression similarity between bulk and pseudo-bulk (PB) profiles, showing overall concordance and transcriptome-wide divergence patterns. (**B**) Histogram of mean expression differences between bulk and PB profiles across all genes, illustrating the distribution and magnitude of bulk–PB shifts across the transcriptome.

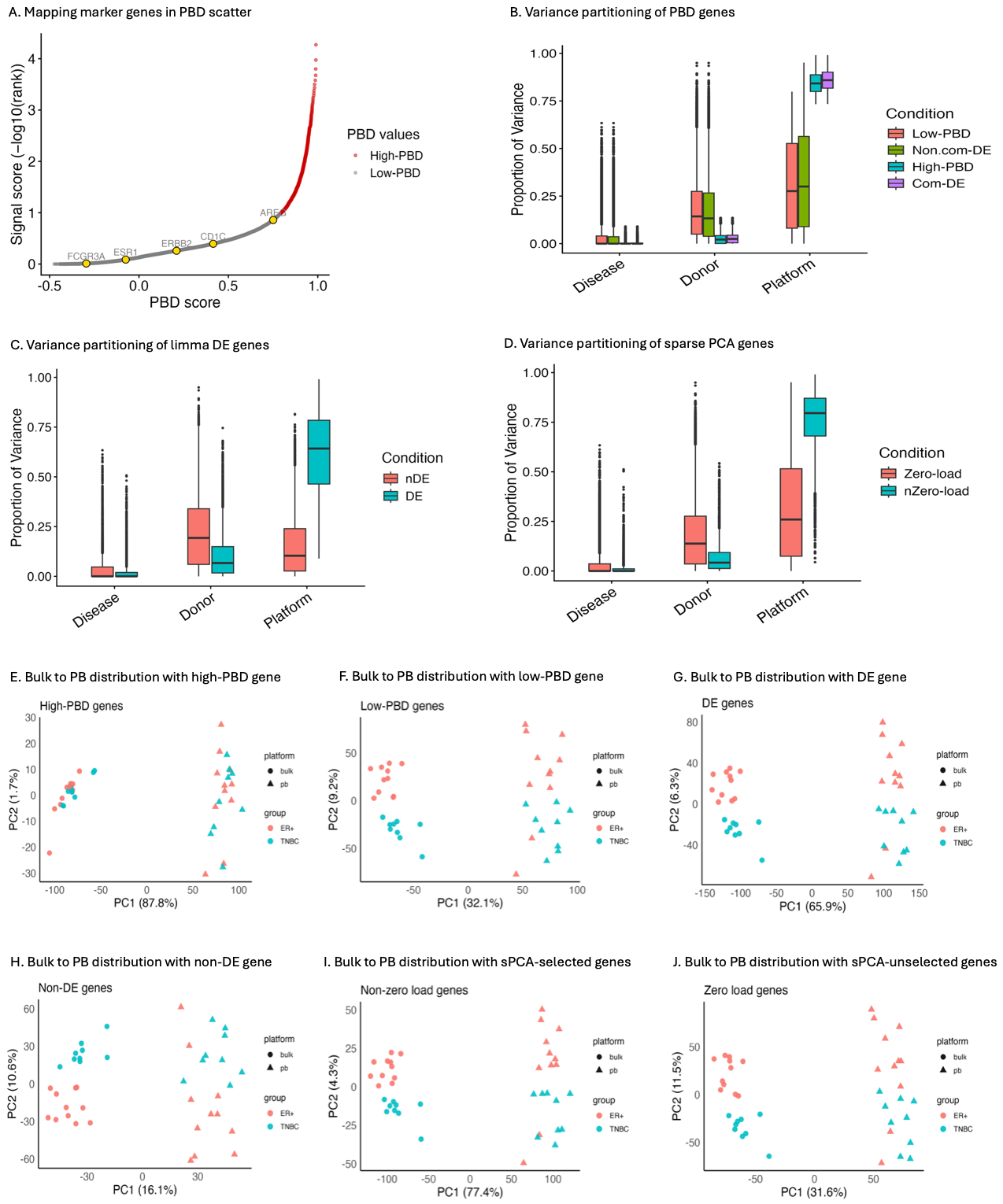

**Supplementary Figure S2. Characterization of PBD-driven variation and bulk–PB distribution patterns in breast cancer data**

(**A**) Marker genes mapped onto the PBD score plot show that high-PBD divergence (red dots) does not eliminate subtype-specific signals. (**B–D**) Variance partitioning of DE genes identified by PBD, limma-DE, and sPCA revealed that high-PBD genes are driven almost exclusively by platform-related effects, with a median of ~98% of their variance explained by platform differences and negligible contributions from disease subtype or donor variation (B). In contrast, genes selected by limma-DE or sPCA capture a mixture of true biological subtype signals and platform-induced biases, demonstrating that these conventional approaches are limited in their ability to disentangle genuine biology from platform-driven variation. (**E–F**) Bulk–PB distributions for high- and low-PBD genes: high-PBD genes show strong divergence between bulk and PB profiles (E), whereas low-PBD genes exhibit reduced platform bias along PC1 but retain subtype separation along PC2 (F). (**G–H**) Bulk–PB distributions for DE and non-DE genes: DE genes display a strong biological signal across platforms and subtypes (G), whereas non-DE genes show confounded divergence across platforms and subtypes (H). (**I–J**) Bulk–PB distributions for sPCA-selected (non-zero loading) and sPCA-unselected (zero loading) genes: sPCA-selected genes capture variance contributed by both subtype and platform effects, while sPCA-unselected genes reflect confounded subtype and platform signals.

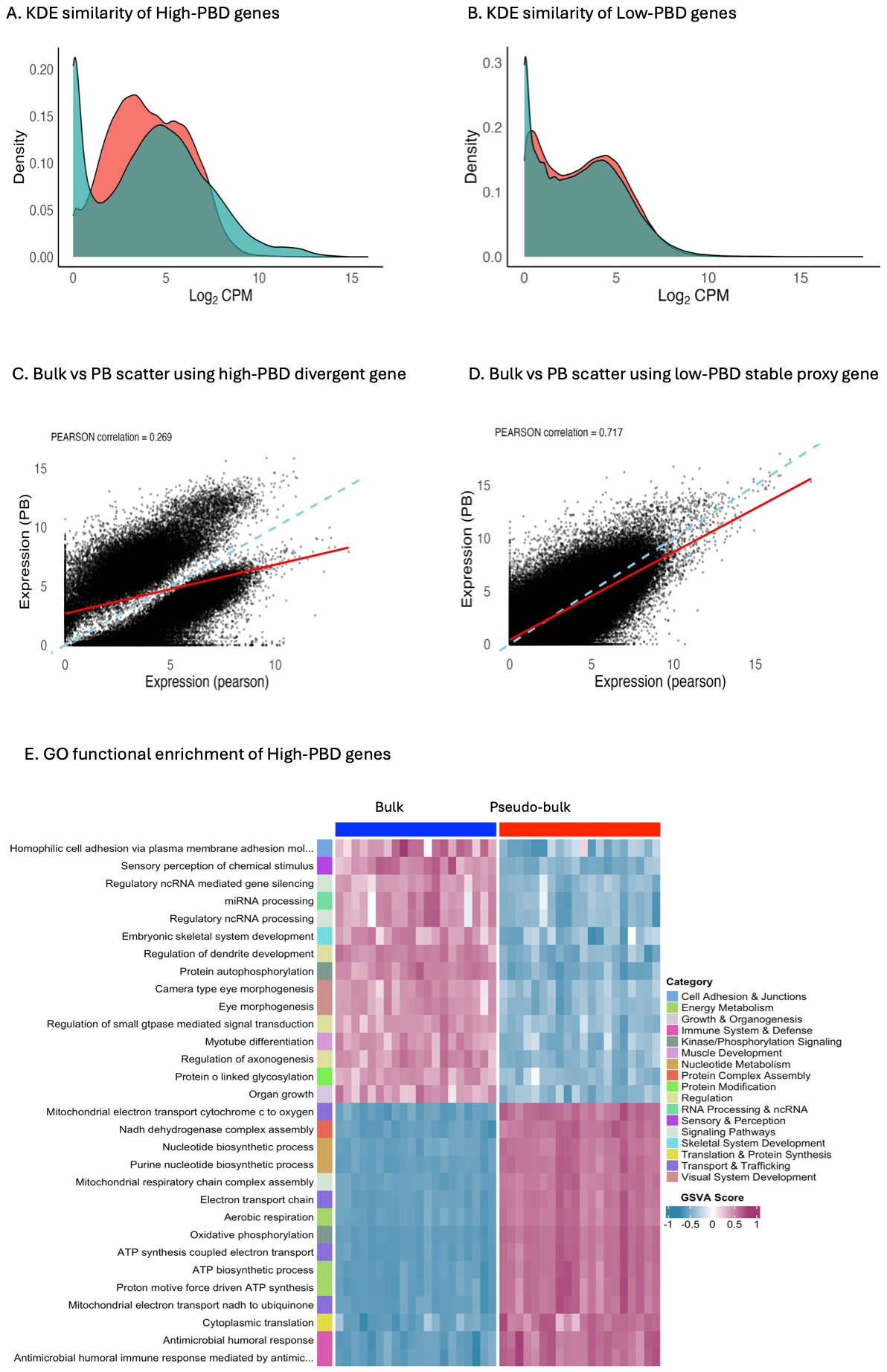

**Supplementary Figure S3. Gene expression similarity and gene ontology (GO) analysis using PBD-stratified genes.**
(**A**) Kernel density estimation (KDE) of bulk–pseudo-bulk expression for high-PBD genes, revealing strong cross-platform divergence. (**B**) KDE of low-PBD genes, demonstrating improved concordance between bulk and pseudo-bulk profiles. (**C-D**) Scatter plots comparing bulk vs. pseudo-bulk expression using: (i) high-PBD drivers, and (ii) low-PBD stable proxy genes, illustrating how PBD stratification influences expression shifts. (**E**) Enriched GO biological processes for high-PBD genes, highlighting pathways associated with platform-driven variability and divergent expression programs.

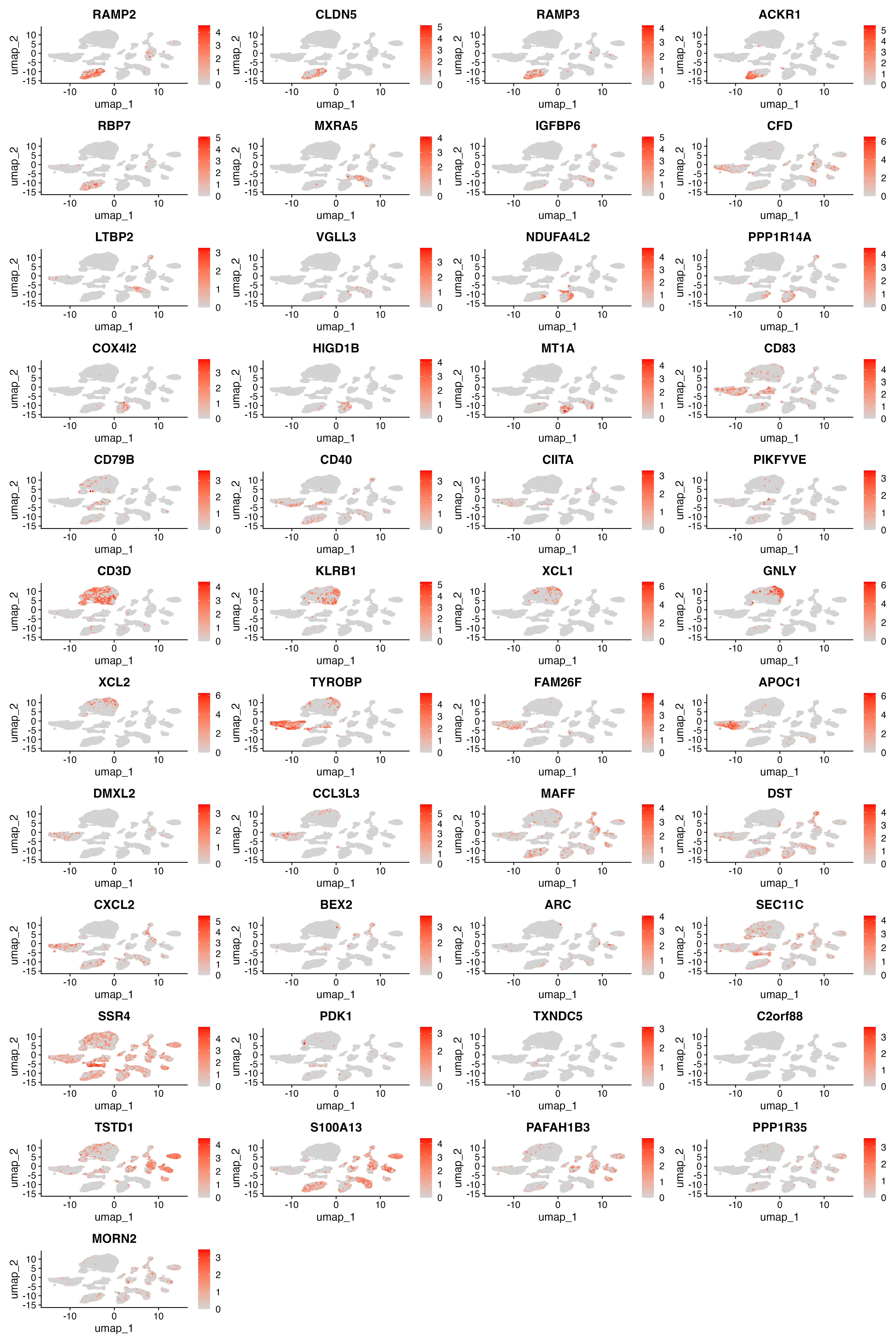

**Supplementary Figure S4A. High-PBD genes exhibit distinct cell–type–associated expression programs**

Projection of cell–type–associated high-PBD genes in the single-cell embedding, showing that highly divergent (high-PBD) genes remain strongly linked to specific cellular populations.

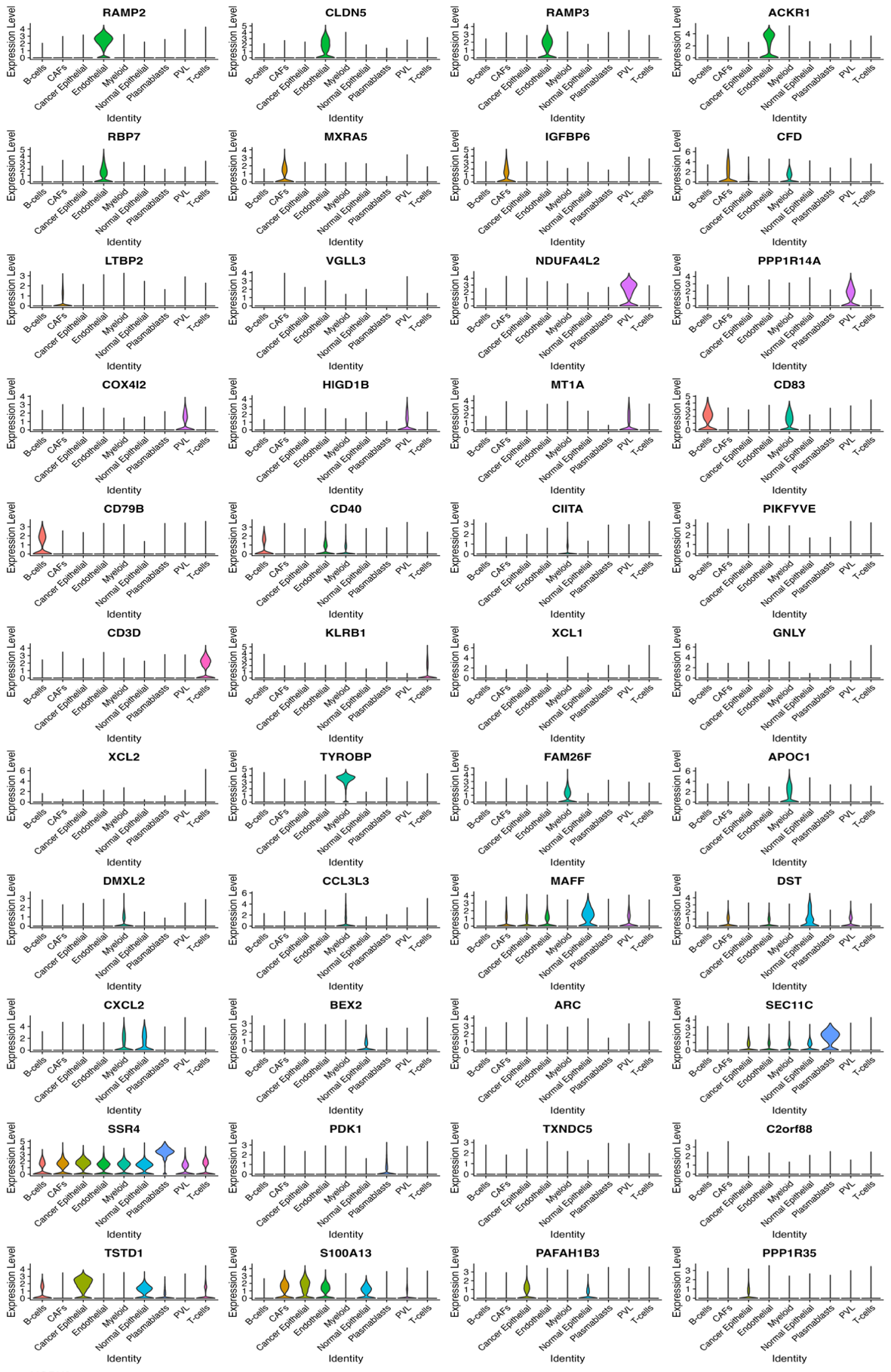

**Supplementary Figure S4B. High-PBD genes exhibit distinct cell–type–associated expression programs**

Violin plots of high-PBD DE genes across annotated single-cell types, illustrating that despite their platform divergence, these genes maintain clear cell–type–specific expression patterns.

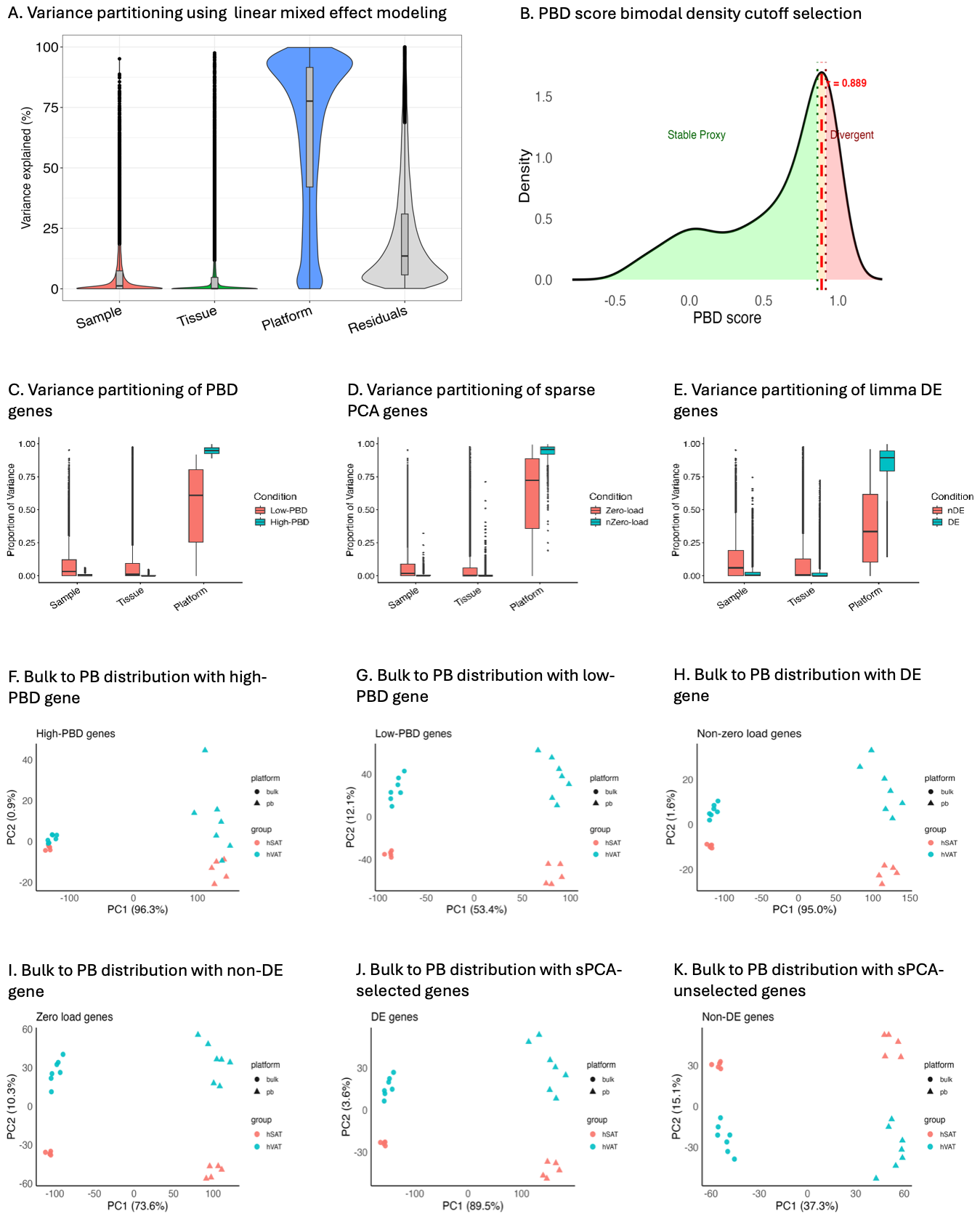

**Supplementary Figure S5. Assessing PBD scores to reveal bulk–pseudo-bulk (PB) distribution discrepancies in human adipose tissue.**

(**A**) Variance decomposition shows the platform explains ~76% of expression variability after adjusting for tissue and sample effects. (**B**) PBD score density with high-PBD cutoff in red, separating cross-platform divergent and stable genes. (**C–E**) High-PBD genes are platform-driven (C); limma DE and sPCA genes are confounded by tissue and sample effects (D–E). (**F–G**) Bulk–PB distributions: high-PBD genes drive platform differences (F); low-PBD genes reduce platform bias along PC1 while retaining tissue separation (G). (**H–I**) DE analysis shows both platform-biased DE and stable non-DE genes preserve tissue signals while limiting platform discrepancies. (**J–K**) Both sPCA-selected and unselected genes capture tissue and platform variance, showing confounded, platform-biased signals that limit their ability to identify cross-platform discrepancies.

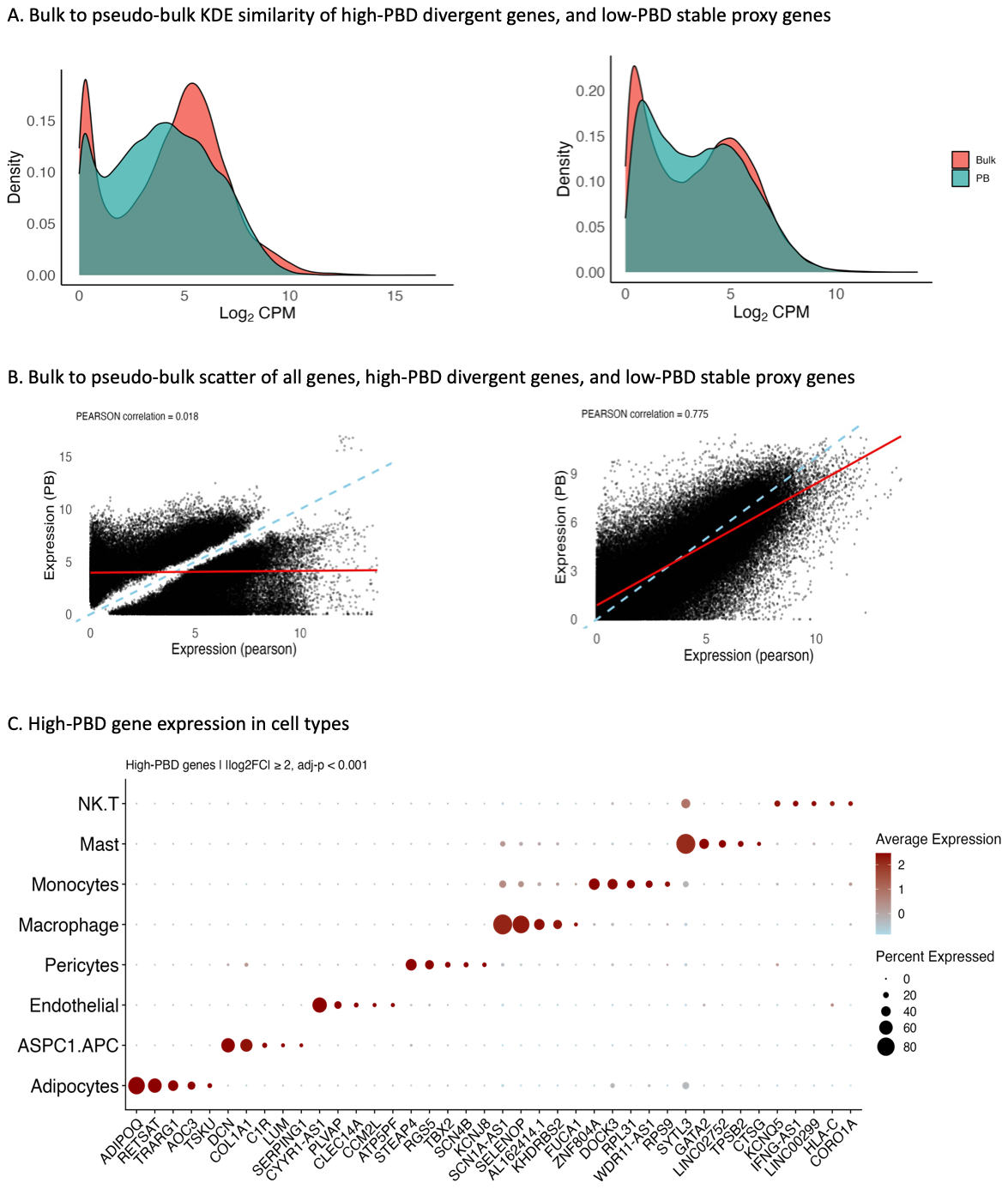

**Supplementary Figure S6. Gene expression similarity comparison using PBD-stratified genes in human adipose tissue.**

(**A**) Kernel density estimation (KDE) of bulk–pseudo-bulk expression for high-PBD genes, showing platform-driven differences and enhanced concordance for low-PBD stable proxy genes. (**B**) Bulk vs. pseudo-bulk scatter plots for high-PBD drivers, and low-PBD stable genes; high-PBD genes show mean shifts, and their removal significantly enhanced cross-platform concordance. (**C**) Expression patterns of high-PBD genes highlight cell–type–specific populations driving cross-platform discrepancies in human subcutaneous (hSAT/SC) tissue.

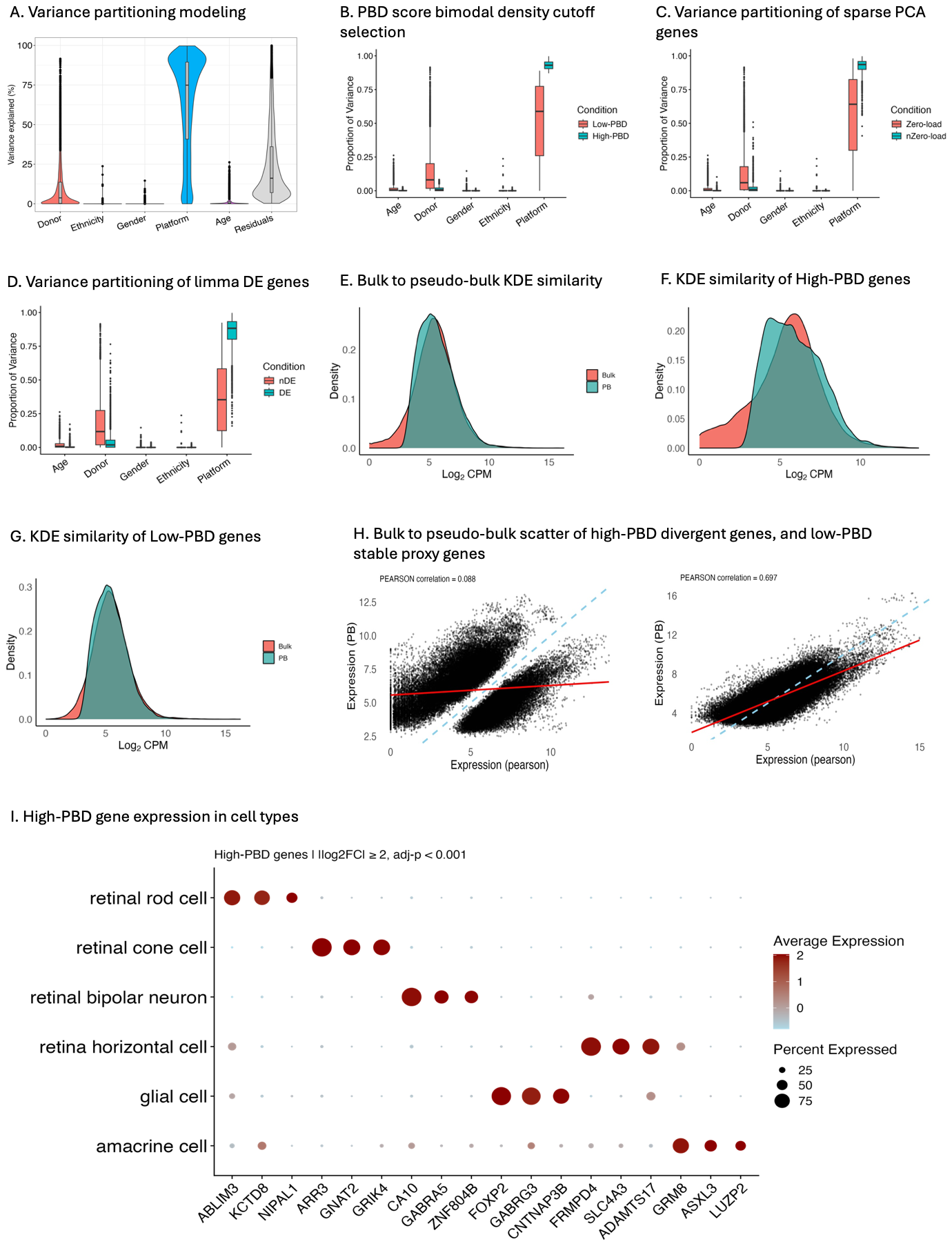

**Supplementary Figure S7. Assessing PBD scores to reveal bulk–pseudo-bulk (PB) discrepancies in human retinal tissue.**

(**A**) Variance decomposition shows that the platform explains a median of 75% expression variability after adjusting for donor and demographic factors. (**B–D**) High-PBD genes are strongly platform-driven, while limma DE and sPCA genes show mixed platform, age, and donor effects. (**E–G**) KDE analyses reveal systemic bulk–PB differences, strongest for high-PBD genes, with improved similarity in low-PBD stable genes. (**H**) Bulk–PB scatter plots reveal that high-PBD genes are major drivers of divergence; removing them significantly enhances cross-platform similarity. (**I**) High-PBD genes exhibit cell–type–specific patterns underlying bulk–PB discrepancies.
